# Exploring the Neural Correlates of Body Perception Disturbance with a Visual Hand Morph Illusion in Complex Regional Pain Syndrome: an fMRI study

**DOI:** 10.1101/2025.10.29.685372

**Authors:** Matthew Mockford, Massieh Moayedi, Jennifer S. Lewis

**Author notes:** **Corresponding Author Contact:** Jennifer S. Lewis, School for Health and Social Wellbeing, Glenside Campus, University of the West of England, Blackberry Hill, Stapleton, Bristol, BS16 1DD, UK.

## Abstract

Complex regional pain syndrome (CRPS) is a debilitating chronic pain condition whose causal mechanisms remain poorly understood. Individuals with CRPS often experience distorted perceptions of their affected limb in terms of size and shape, accompanied by feelings of dislike and disownership— collectively referred to as body perception disturbances (BPD). Evidence that both BPD and pain can rapidly reduce following a visual hand morph illusion suggests modulation of central body representations. To elucidate the neural correlates underlying these rapid perceptual changes, we investigated brain activity during the hand morph illusion using functional magnetic resonance imaging (fMRI).

Forty-four participants (22 with upper limb CRPS, 22 pain-free controls) completed the hand morph illusion task during fMRI and provided trial-by-trial ratings of ownership, likeability, and pain of their affected hand.

People with CRPS showed greater activation of the left premotor cortex (PMC) during the illusion compared with controls. Functional connectivity between the left PMC and right extrastriate body area (EBA) was also revealed in CRPS during the illusion. No main effects of the illusion, relative to a sham condition, were found on pain intensity, limb ownership, or likeability ratings.

These findings provide the first identification of neural correlates of body perception disturbance in CRPS, implicating PMC–EBA networks in altered body representation. Further neuroimaging research is required to define the neural signature underlying this disturbance.

## Introduction

Complex regional pain syndrome (CRPS) is a debilitating primary pain condition which can affect a limb after a peripheral noxious event such as surgery, fracture or trauma[8]. CRPS is characterized by severe pain, swelling, colour changes, and motor difficulties[3; 26]. While the etiology of CRPS is poorly understood, evidence indicates that changes in the peripheral and central nervous system (CNS) contribute to its pathophysiology[11]. Prospective studies suggest that 27% of acute CRPS cases progress to persistent symptoms, leading to significant functional impairment and negative impact on quality of life[2; 21; 33]. Given the significant individual and economic burden posted by CRPS, developing effective treatments that target underlying mechanisms of the disease are needed[36].

People with CRPS report body perception disturbances (BPD) of their affected limb: disowner ship and inattention of the limb[23], distorted body image (size, shape, or colour)[25], and a blurred representation of position and movement of the painful limb[4]. As such, people with CRPS have an altered perception of their affected limb while the rest of their body is perceived as normal[1]. BPD can be measured by the clinician-administered and validated Bath BPD scale which has utility as a modifiable measure[23]. Those who report more extensive disturbances in limb perception also rate greater pain intensity, and multidisciplinary rehabilitation leads to reductions in BPD, suggesting a relationship between pain and bodily representations[22; 25].

Multisensory bodily illusions can modify central body representations such as body image, body schema, and limb ownership[5; 29]. The MIRAGE system uses augmented reality to create illusions which manipulate perception of the painful limb such that body representations are modulated in real-time[30]. Using the MIRAGE system, a visual hand morph illusion reduced BPD and pain intensity in individuals with CRPS[24]. However, the neural correlates of this analgesic illusion are unknown. We have previously shown using functional magnetic resonance imaging (fMRI) that a network of brain regions including the extrastriate body area (EBA), posterior parietal cortex (PPC), and left ventral premotor cortex (vPMC) are responsible for changes in body image in healthy individuals using a multisensory finger stretch illusion in the MIRAGE system[27]. Given the role of BPD in CRPS, it is feasible that activity in these regions during bodily illusion will differ from pain-free individuals.

This study aims to explore the analgesic effect of the hand morph illusion by capturing neural activity in response to a modified illusion performed within an MRI environment[27]. This allows for the identification of brain regions and mechanisms which may mediate the relationship between body representations and pain in individuals with CRPS. We hypothesize that brain activity in regions involved in body image (EBA), body schema (PMC, PPC), and pain modulation (insula, anterior cingulate cortex, primary somatosensory cortex) will differ in those with CRPS compared to pain-free individuals. Uncovering the neural mechanisms of BPD will allow for a better understanding of CRPS pathophysiology, leading to improved treatments for individuals living with this debilitating condition.

## Methods

### Participants

Twenty-eight participants who met the Budapest criteria [16] for CRPS affecting one upper limb were recruited from the National CRPS Service, Royal United Hospital NHS Trust Bath or CRPS UK network registry (crpsnetworkuk.org/Registry). Additionally, 29 healthy, pain-free individuals were recruited as age-matched controls.

The study was approved by South Central Research Ethics Committee (REC reference: 17/SC/0649), NHS Health Research Authority (IRAS Project ID: 234938) and University Ethics Committees. It was conducted in accordance with the Declaration of Helsinki All participants provided written informed consent prior to their inclusion in the study.

Ten participants (five with CRPS, five who are pain-free) only completed the behavioural session and did not undergo the neuroimaging session, one participant was missing a structural T1-image and registration could therefore not be completed, and three participants had excessive head motion (>3mm) during the task fMRI scan. This resulted in a final sample of 22 individuals with CRPS (two males; 20 females; aged 52.77 ± 11.3) and 22 pain-free controls (five males, 17 females, aged 53.09 ± 14.43).

### Experimental Design

#### Behavioural Session

Data collection occurred over two visits up to two weeks apart. The first visit was a behavioural session where the hand morph illusion was calibrated (followed by a second fMRI session explained below). This session took place at the Wolfson Centre Research Facility in Bath, United Kingdom. After providing informed consent, participants completed questionnaires, filled in painful regions on a body map (to confirm controls were pain free). The Bath Body Perception Disturbance Scale [24] and the CRPS Symptom Severity Score (CSS) [17]were administered by JL.

Individuals with CRPS viewed their affected limb on a screen inside the MIRAGE system where two distinct images of the hand were created: 1) an image of the hand without any manipulation (’REAL’) and 2) a desired version of the hand which is how they would like it to look (‘DESIRED’). The MIRAGE system operator digitally altered the appearance of the painful hand in response to the specific description given by each participant based on how they wished their hand to look. visual changes to aspects of shape, size and/or colour of the hand, were digitally altered in real time to create the hand illusion. (see Lewis et al 2021 for further detail about the procedure). A digital illusion template with these specific parameters was saved for each participant to re-create the hand illusion in the scanner

Pain free controls: the same MIRAGE procedure was performed as previously described using the participant’s dominant hand apart from the hand illusion comprised visually enlarging the hand image by a 30% in overall size to create the ‘Distorted’ hand illusion template.

In preparation for the MRI session, all participants performed 4 practice trials of the illusion. Examples of hand morph illusion can be seen in Figure 1. The practice runs of the illusion were presented on a monitor at eye level to mimic the setup inside the MRI scanner.

**Figure 1:**
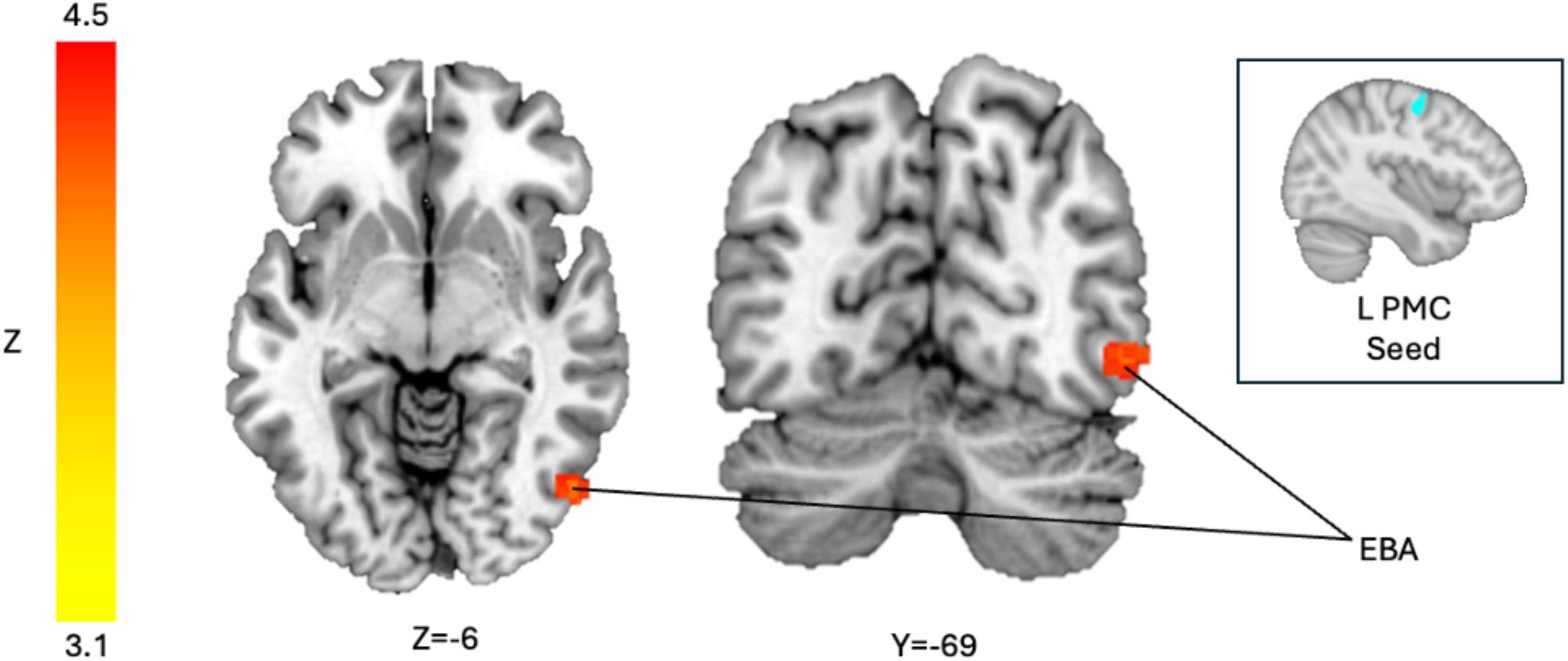
Hand Morph Illusion. Examples of a hand morph illusion in three participants with CRPS. The panels on the left represent the actual, unmanipulated hand, and the panels on the right represent the desired hand as described by those living with CRPS.

#### Neuroimaging Session

The neuroimaging session took place at the Centre for Integrative Neuroscience and Neurodynamics, University of Reading, Berkshire, UK using 3T Siemens PrismaFIT MRI scanner (Erlangen, Germany) using a 64-channel head/neck coil. Before entering the scanner participants had vac-loc cushions to support the experimental arm in position. The MRI session comprised five scan protocols: an anatomical T1-weight sequence, resting state arterial spin labelling, and a resting-state fMRI, and a hand morph illusion fMRI task. Participants were presented with 40 trials (across four runs, ten trials per run), 20 of which were the hand morph illusion, and 20 control trials with a static hand image, presented in a pseudo-randomized order. The illusion was viewed upon an MRI compatible LCD screen (BOLDScreen) via a mirror mounted on the attached to the head coil. For each trial of the illusion (See Figure 2), participants with CRPS were presented with their unmanipulated ‘REAL’ hand (3 s) changing into the previously calibrated ‘DESIRED’ image of their hand (‘morph’ phase of the illusion). They then looked at the ‘DESIRED’ image of the hand for 3 seconds (‘hold’ phase of the illusion) before it changed back to the unmanipulated image. The control image consisted of an unmanipulated image of their ‘REAL’ hand for 9 seconds while a black bar moved across the image to control for observing movement. Healthy controls underwent a similar illusion protocol, however the ‘morph’ phase of the illusion displayed the ‘DISTORTED’ version of their hand changing into an image of their ‘REAL’ hand, before going back to the ‘DISTORTED’ image. For control trials of healthy volunteers, the image stayed as the ‘DISTORTED’ hand image with a black bar moving across the image. After a 4-8 second jitter period, participants rated their current hand pain, how much they believe the hand they saw belonged to them, and how much they like the appearance of the hand they saw. Participants were given 5 seconds to report their ratings on a visual analog scale (VAS) from 0 to 100. Each run was 6 minutes and 37.5 seconds with a 2-minute break between runs.

**Figure 2:**
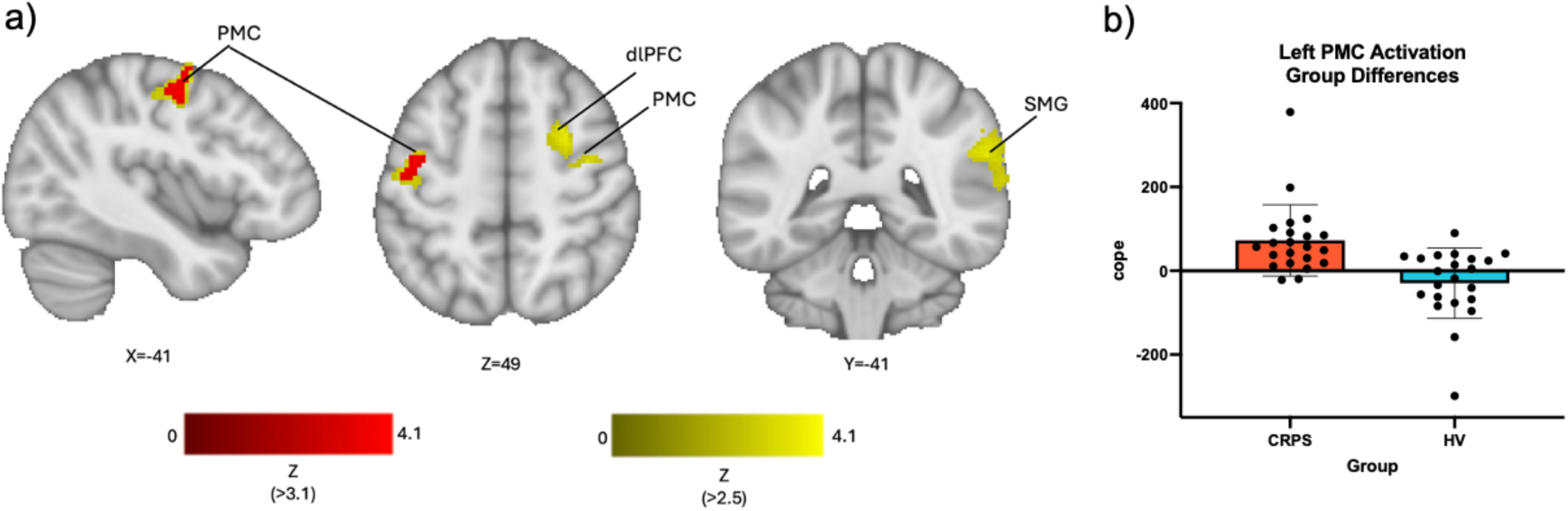
Differences in brain activation during the hand morph illusion in CRPS versus pain-free participants. a) Statistical images are cluster-corrected pFWE<0.05 (cluster-forming height threshold Z>3.1 [red] or Z>2.5 [11]). b) Group differences in contrast parameter estimates of left premotor cortex (PMC). Abbreviations: PMC—premotor cortex; dlPFC—dorsolateral prefrontal cortex; SMG— supramarginal gyrus.

### Functional Magnetic Resonance Imaging Acquisition and Analysis

#### Acquisition Parameters

Five scan protocols were collected in total, however there are two relevant to the current study. The first, an anatomical three-dimensional magnetization-prepared, rapid acquisition gradient echo (MPRAGE) scan (TR = 2,300 s, TE = 2.29 ms, slice thickness = 0.94 mm, voxel size = 0.94 mm^3^). The second, an fMRI scan was collected during the hand illusion, with the following parameters: 405 volumes acquired using a T2*-weighted echo-planar imaging sequence (TR = 1000 ms, TE = 30 ms, flip angle = 90°, slice thickness = 2mm, voxel size= 2 x 2 x 2 mm^3^). Participants underwent four runs of the task. Other scan types collected, but not included herein, are include a resting state fMRI, and an arterial spin labeling resting state scan.

#### Preprocessing

Raw DICOM data converted and curated according to the Brain Imaging Data Structure (BIDS, https://bids.neuroimaging.io) with dcm2bids v.2.1.4 (dcm2niix v.1.0.20201102). Results included in this manuscript come from preprocessing performed using fMRIPrep 24.0.1 [9; 10] which is based on Nipype 1.8.6[13].

Anatomical T1*-weighted images were corrected for intensity non-uniformity with N4BiasFieldCorrection[37], and skull-tripped with ANTs (v2.5.1). Tissue segmentation completed using FSL’s FAST software (v5) and surface based reconstructed with Freesurfer’s recon-all (v7.3.2) [7; 42]. Normalization to standard space using nonlinear registration performed with ANTs (v2.5.1) with templates accessed from TemplateFlow (v24.2.0)[6].

For the illusion task functional scan, BOLD references were co-registered to the T1w references using Freesurfer’s bbregister (v7.3.2)[14]. Preprocessed functional volumes were denoised using independent component analysis (ICA) in FSL’s FIX. Components were classified manually using MELODIC (Multivariate Exploratory Linear Optimized Decomposition into Independent Components[15; 35]. Finally, the denoised data was spatially smoothed using a 6mm (FWHM) Gaussian kernel with FSL.

#### fMRI Hand Morph Illusion Analysis

After preprocessing, fMRI analysis was carried out using FMRI Expert Analysis Tool (FEAT) Version 6.00, part of FMRIB’s Software Library (FSL; www.fmrib.ox.ac.uk/fsl). A three-level general linear model (GLM) was carried out on the final preprocessed data. First level statistical analysis was performed modeling each phase of the illusion as a separate regressor (active morph, sham morph, active hold, sham hold, rating). Some participants had large holes in their brain mask after first-level analysis, so we used Synthstrip (v1) to further remove non-brain voxels and correct brain masks[18]. Second-level fixed-effect analysis were utilized to generate contrast of parameter estimate (COPE) images to produce an average image for each participant. Third-level mixed-effect analysis using FMRIB’s Local Analysis of Mixed Effects-1+2 (FLAME-1+2) produced an average group map image for the CRPS group and the pain-free group where the following contrasts were modelled to compare brain activity between groups: CRPS group average, pain-free group average, CRPS>pain-free, and pain-free>CRPS. Z (Gaussianised T/F) statistic images were thresholded using clusters determined by Z>3.1 and a corrected cluster significance threshold of p=0.05[39]. Sex was included as a nuisance regressor. We also ran the group difference analysis with a more liberal cluster-forming threshold of Z > 2.5.

#### Psychophysiological Interaction

Lastly, we completed a psychophysiological interaction (PPI) analysis within the CRPS group to investigate task-based functional connectivity of brain regions identified in the hand morph task analysis[31]. ROI seeds were selected by the peak activation from the CRPS>control contrast, specifically the left PMC.

The PPI analysis was performed by modelling the interaction of the extracted time-series of the ROI (physiological regressor), and the onset of the hand morph (psychological regressor) in a GLM. Second-level participant average contrasts of the PPI were created using fixed-effect analysis. At the group-level, a mixed effect analysis (FLAME 1+2) was used to compare connectivity maps within the CRPS group. Statistical images were thresholded using clusters determined by Z>3.1 and a corrected cluster significance threshold of p=0.05. Sex was included as a nuisance regressor.

## Results

Key demographic and behavioural information for the two groups can be found in Table 1 for the CRPS group and Table 2 for the pain-free controls. The questionnaires were intended to evaluate clinical factors in CRPS (i.e., BPD and CSS), so were not compared across groups. As the primary goal of this study was to investigates brain activity, the ratings collected in the scanner will not be reported here, but can be found in the Supplementary Materials.

**Table 1:**
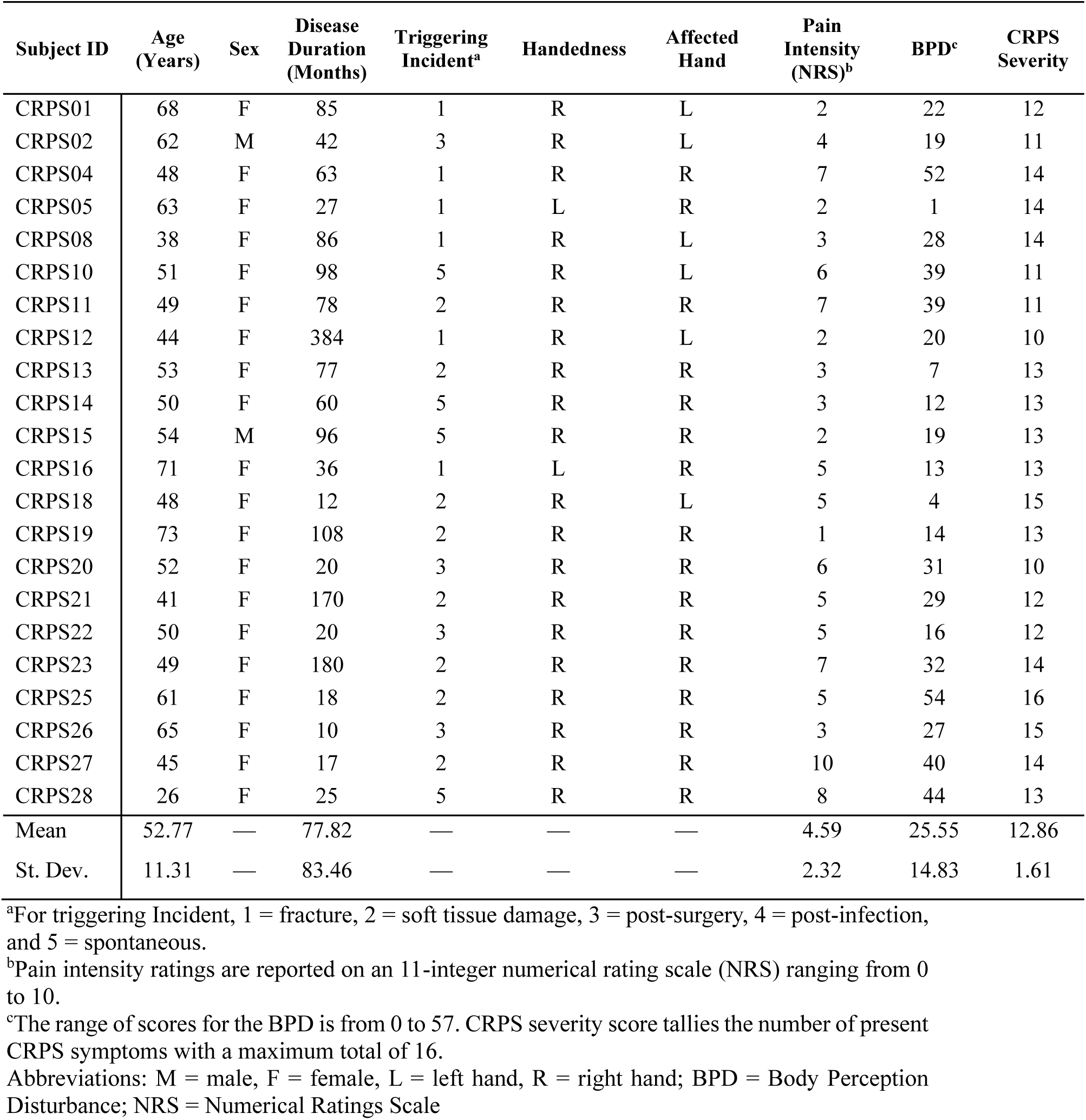
CRPS Participant Information.

**Table 2:**
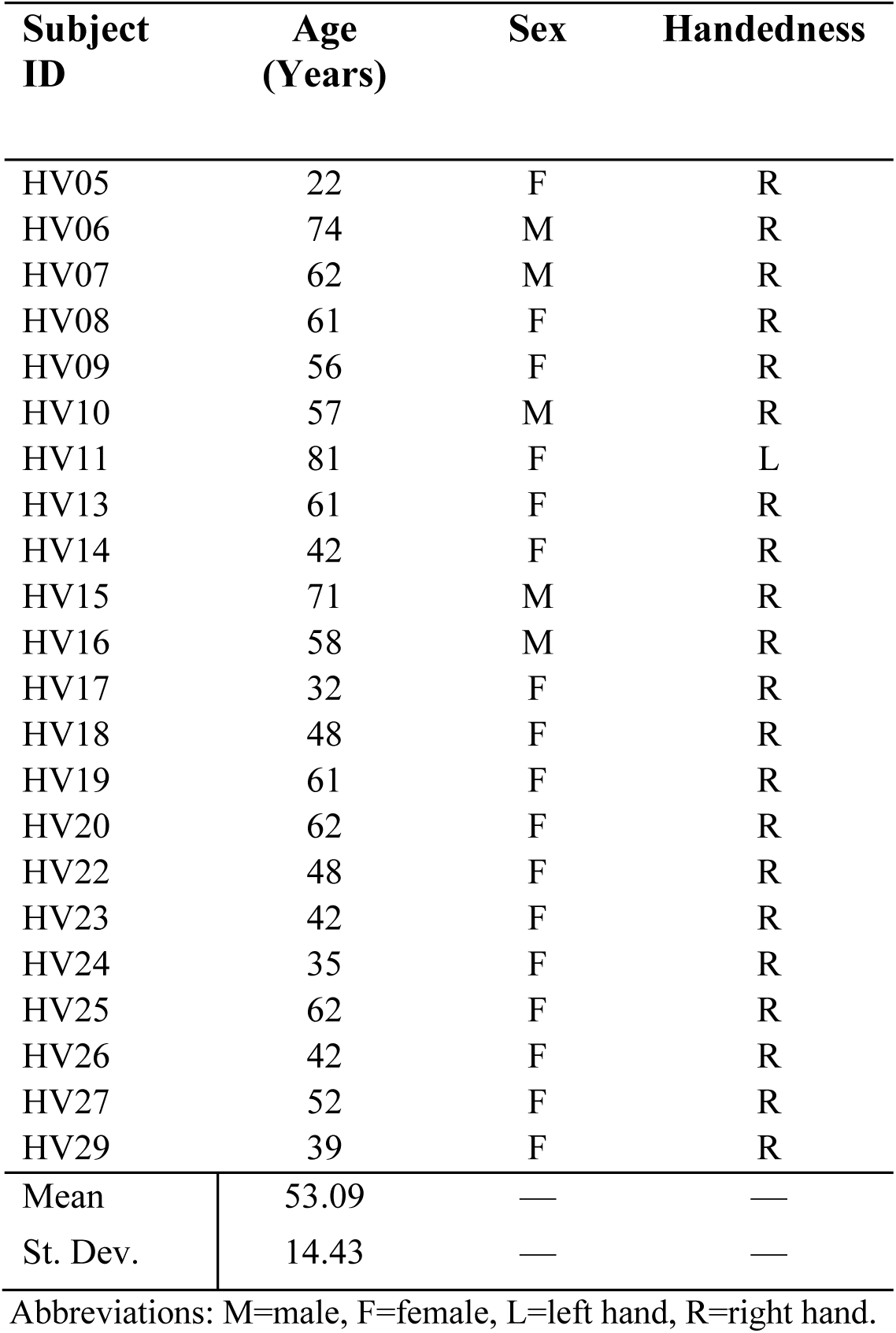
Pain-Free Control Participant Information.

| **Subject ID** | **Age (Years)** | **Sex** | **Handedness** |
| --- | --- | --- | --- |
| HV05 | 22 | F | R |
| HV06 | 74 | M | R |
| HV07 | 62 | M | R |
| HV08 | 61 | F | R |
| HV09 | 56 | F | R |
| HV10 | 57 | M | R |
| HV11 | 81 | F | L |
| HV13 | 61 | F | R |
| HV14 | 42 | F | R |
| HV15 | 71 | M | R |
| HV16 | 58 | M | R |
| HV17 | 32 | F | R |
| HV18 | 48 | F | R |
| HV19 | 61 | F | R |
| HV20 | 62 | F | R |
| HV22 | 48 | F | R |
| HV23 | 42 | F | R |
| HV24 | 35 | F | R |
| HV25 | 62 | F | R |
| HV26 | 42 | F | R |
| HV27 | 52 | F | R |
| HV29 | 39 | F | R |
| Mean | 53.09 | — | — |
| St. Dev. | 14.43 | — | — |

Abbreviations: M=male, F=female, L=left hand, R=right hand.

To determine whole brain group-level activations we modeled the ‘hold’ phase of the hand morph illusion where the participant viewed the final result of the illusion. Thus, any referencing to the effect of the illusion moving forward specifically refers to this portion of the illusion. The ‘morph’ phase where the hand actively changes appearance did not yield any significant group level results. We determined whether brain regions had differing activity between the CRPS group and pain-free group during the illusion. At a cluster-forming threshold of 3.1, we discovered that the left PMC had greater activation during the illusion in people with CRPS than in pain-free individuals. When we conducted the same analysis with a more liberal cluster-forming threshold, the bilateral PMC, right dorsolateral prefrontal cortex (dlPFC) and right supramarginal gyrus had greater activation during the illusion in those with CRPS than in pain-free individuals. These findings are summarized in Figure 2 and Table 3.

**Table 3:** Brain Activity differences during hand morph illusion between CRPS and pain-free groups.

| Region | Z score | Peak MNI |  |  | Cluster Extent<br>(Voxels) |
| --- | --- | --- | --- | --- | --- |
|  |  | X | Y | Z |  |
| Cluster-forming height threshold: Z > 3.1, extent threshold p=0.05 |  |  |  |  |  |
| L PMC | 4.1 | -36 | -4 | 54 | 133 |
| L PMC | 4.03 | -42 | -10 | 54 |  |
| L PMC | 3.97 | -42 | -8 | 50 |  |
| L PMC | 3.73 | -48 | -14 | 50 |  |
| L PMC | 3.61 | -42 | -4 | 60 |  |
| Cluster-forming height threshold: Z > 2.5, extent threshold p=0.05 |  |  |  |  |  |
| R SMG | 3.97 | 60 | -42 | 26 | 367 |
| R SMG | 3.59 | 60 | -40 | 30 |  |
| R SMG | 3.44 | 62 | -36 | 28 |  |
| R SMG | 3.43 | 54 | -42 | 28 |  |
| R SMG | 3.43 | 64 | -46 | 16 |  |
| R SMG | 3.3 | 64 | -42 | 24 |  |
| R PMC | 4.4 | 48 | -4 | 56 | 343 |
| R PMC | 3.88 | 38 | -8 | 46 |  |
| Outside BAs | 3.87 | 34 | -16 | 38 |  |
| R SMA | 3.81 | 26 | 2 | 48 |  |
| Outside BAs | 3.37 | 40 | -12 | 32 |  |
| R PMC | 3.36 | 38 | -12 | 44 |  |
| L PMC | 4.16 | -38 | -6 | 54 | 325 |
| L M1 | 3.71 | -46 | -14 | 52 |  |
| L PMC | 3.47 | -42 | -8 | 46 |  |
| L M1 | 3.37 | -46 | -12 | 36 |  |
| L PMC | 3.33 | -42 | -4 | 60 |  |
| L M1 | 3.15 | -48 | -14 | 40 |  |

Statistical threshold was set cluster-corrected $pFWE < 0.05$ (cluster-forming height threshold $Z > 3.1$ and $Z > 2.5$ ).
MNI coordinates refer to local maxima from each cluster.
Abbreviations: L = left, R = right. PMC = premotor cortex, M1 = primary motor cortex, SMG = supramarginal gyrus

Next, we ran a PPI analysis within the CRPS group to see whether there was task-based functional connectivity of the left PMC with the rest of the brain. We discovered that in individuals with CRPS, the left PMC is significantly functionally connected to the right EBA during the illusion (Figure 3).

**Figure 3:**
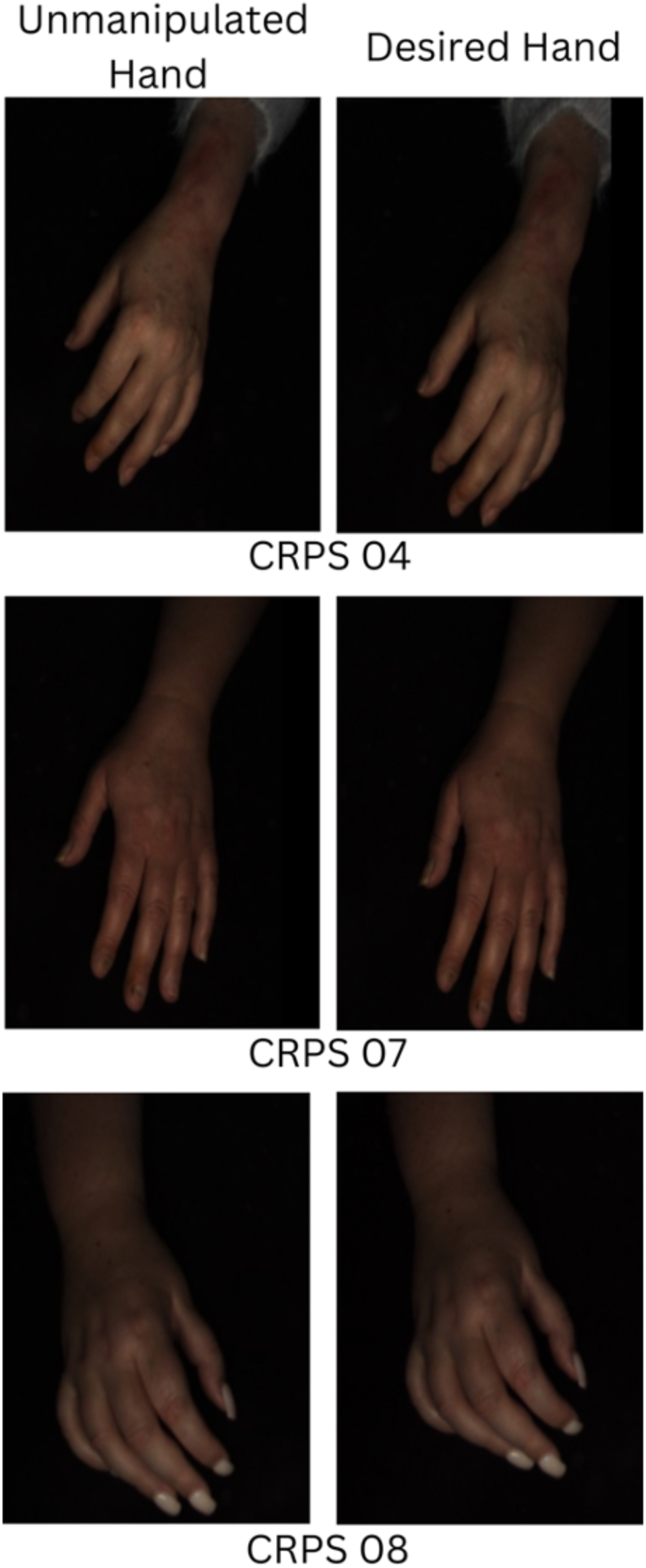
Task-Based Functional Connectivity of Left Premotor Cortex in CRPS. Psychophysiological interaction (PPI) analysis of the left premotor cortex (PMC) revealed significantly connectivity to the right extrastriate body area (EBA) in CRPS. Statistical images are cluster-corrected p<0.05 (cluster forming threshold Z>3.1).

## Discussion

This study aimed to identify the neural underpinnings of body perception disturbance in people with CRPS via an analgesic hand morph illusion[24]. To investigate this, we compared activity in the brain between individuals with CRPS and pain-free controls. We discovered that individuals with CRPS had greater activity in the bilateral PMC, right dlPFC, and right SMG. We further reported that within the CRPS group, the left PMC was functionally connected to the right EBA during the illusion. Given the role of BPD in CRPS, we predict that these regions may play a role in maintaining pain symptoms by disturbing perceptions of the painful limb.

Individuals with CRPS had greater activity in the left PMC compared to pain-free controls during the hand morph illusion. One study has previously implicated the PMC in CRPS symptomology where participants with dystonia had reduced activity in the PMC ipsilateral to the affected limb during a motor imagery task compared to healthy controls[12]. In our study, we observed greater PMC activity in individuals with CRPS during the illusion. However, given the differences between the tasks and CRPS symptomology between the two groups, it is possible that the PMC plays more than one role in the pathophysiology of CRPS. While its name implies the PMC’s role in the preparation of motor activity, its role in limb ownership and embodiment has been demonstrated using the rubber hand illusion (RHI)[34; 41]. More specifically, the ventral PMC has been implicated in dynamic limb ownership, as demonstrated by the lack of embodiment in the RHI in lesion patients[41]. Alternatively, the dorsal PMC is responsible for receiving information from different sensory modalities and using it to prepare for goal-oriented motor behaviour (which relies on the body schema—a map of the body in space)[19; 28]. Given the PMC’s role in limb ownership and body schema, it is well posited to play a multi-faceted role in CRPS pathophysiology.

Given the relatively small sample size, we lowered the cluster-forming threshold to a more liberal level of Z > 2.5, we found that the right dlPFC also had greater activity in individuals with CRPS than pain-free controls during the illusion. It has been proposed that the dlPFC is involved in storing sensory information pertaining to peri-personal space so that it can be used to plan motor behaviour[19]. Studies that have attempted to differentiate the role of the dlPFC and PMC in motor behaviour by splitting tasks into progressive stages of perception and action suggest that the dlPFC remains active while maintaining the spatial location of cues, while the dorsal PMC encodes finer features of movement such as direction and angle [40] Given that this cluster overlaps both right PMC and dlPFC, it is possible that both regions are playing a similar role in the illusory limb’s incorporation of the body schema.

The serial co-construction model of body representations has been used to explain how ameliorating disturbances in the perception of the limb can lead to reductions in pain[24]. In this model, body image and body schema are seen as malleable and influencing one another[32]. We have previously proposed that pain in CRPS can be maintained by an incongruence between body image and body schema. By rapidly modulating body image so that the hand becomes a visually desired version, the incongruence is reconfigured, and pain is suppressed indirectly by addressing disturbances in bodily perceptions[24].

In our PPI analysis, we discovered that within the CRPS group the left PMC was significantly functionally connected to the right EBA during the hand morph illusion. Given the role of the EBA in encoding and integrating visual features of body parts (body image) [38]and the PMC in planning and executing motor behaviour (body schema)[40], It is possible that functional connectivity between the EBA and PMC may mediate the incongruencies between body image and schema, positing them as key regions in the brain mechanisms underlying illusory analgesia[24].

The primary goal of this study was to investigate brain regions that may play a role in body perception disturbance with an analgesic hand morph illusion. Certain modifications to the illusion were required for it to be completed within an MRI environment which may have resulted in our difficulties replicating our previous behavioural results. Given the presence of hyperalgesia and allodynia, participants with CRPS are likely to experience discomfort or pain from the positioning of their painful limb in keeping it outstretched and still within the cramped scanner space for the duration of the scan[20]. In addition, participants only viewed the final image of the illusion for 3 seconds in the scanner as opposed to a full 60 seconds in the MIRAGE device in our previous behavioural study[24]. It is possible that for illusory analgesia to occur, the participant requires a longer exposure to the desired hand. In addition, the pseudo-randomized illusion and sham control trial design may limit any cumulative and lasting illusory effects by potential carry-over from preceding trials, making it difficult to accurately measure the intended effect. These limitations may explain the absence of pain reduction in this study, in contrast to the findings from our previous clinical trial [25].

In conclusion, we are the first to identify brain regions associated with body perception disturbance in CRPS. These regions may play a role in mediating the relationship between pain and body representations of the CRPS limb. Our results show that the left PMC and right EBA could play a role in normalizing discrepancies between body image and body schema in individuals with CRPS, supporting Lewis’ co-construction model of illusory analgesia[24]. Understanding the brain regions involved in body perception disturbance opens new targeted treatment avenues in CRPS, as modifying body perception disturbance can lead to meaningful pain reduction for people living with CRPS.

## Acknowledgements

The authors have no conflict of interests to declare. De-identified aggregate data are available upon reasonable request to the corresponding author. This study was made possible by funding by a Versus Arthritis Pain Challenge Research Award (Grant Number: 201503).

M Moayedi is supported by a Canada Research Chair in Pain NeuroImaging, and by the Bertha Rosenstadt Fund at the University of Toronto’s Faculty of Dentistry, and a Pain Scientist Award from the University of Toronto’s Centre for the Study of Pain. M Mockford was supported by a Queen Elizabeth II Graduate Scholarship in Science and Technology (Ontario Government).

## Data Availability Statement

De-identified/Aggregate data are available upon reasonable request to corresponding author.

## Ethics approval Statement

Procedures reviewed and approved by the University of University of the West of England, the University of Reading and the UK National Institutes of Health Research (NIHR) ethics committees.

